# Going through the motions: incorporating movement analyses into disease research

**DOI:** 10.1101/237891

**Authors:** Eric R. Dougherty, Dana P. Seidel, Colin J. Carlson, Orr Spiegel, Wayne M. Getz

## Abstract

Though epidemiology dates back to the 1700s, most mathematical representations of epidemics still use transmission rates averaged at the population scale, especially for wildlife diseases. In simplifying the contact process, we ignore the heterogeneities in host movements that complicate the real world, and overlook their impact on spatiotemporal patterns of disease burden. Movement ecology offers a set of tools that help unpack the transmission process, letting researchers more accurately model how animals within a population interact and spread pathogens. Analytical techniques from this growing field can also help expose the reverse process: how infection impacts movement behaviors, and therefore other ecological processes like feeding, reproduction, and dispersal. Here, we synthesize the contributions of movement ecology in disease research, with a particular focus on studies that have successfully used movement-based methods to quantify individual heterogeneity in exposure and transmission risk. Throughout, we highlight the rapid growth of both disease and movement ecology, and comment on promising but unexplored avenues for research at their overlap. Ultimately, we suggest, including movement empowers ecologists to pose new questions expanding our understanding of host-pathogen dynamics, and improving our predictive capacity for wildlife and even human diseases.

## Introduction

Disease ecology is a fairly young field, especially compared to epidemiology, which dates back centuries. The two fields overlap often, and share a similar goal: to understand, predict, and (sometimes) prevent disease outbreaks. However, disease ecologists face at least two additional challenges unique to wildlife research. First, disease ecology frequently requires a broad, multispecies perspective that captures complex and counter-intuitive ecosystem dynamics; for example, invasive Burmese pythons’ selective feeding within mammal communities has indirectly increased mosquitoes’ feeding on rodents, in turn amplifying the Everglades virus, which causes encephalitis in humans (Hoyer *et al.* 2017). Second, and equally challenging, is the fact that behavior is just as important for wildlife as for human disease, but harder for researchers to directly interrogate. Epidemiologists frequently use interviews and observational work to study how human behaviors such as sexual activity, international travel, or outdoor labor become risk factors for infectious disease—often directly inspiring interventions; animal behavior, while just as important to disease transmission, is harder to observe and predict in nature.

Movement ecology, also a comparatively young field, uses high-resolution spatiotemporal data to make sense of animal behavior. The “movement ecology paradigm” treats movement as the outcome of behavioral decisions influenced by the interplay of animals’ internal states (e.g., physiological needs), external biological factors (e.g., predation or competition), and the physical environment (e.g., mountain ranges or water sources) (Nathan *et al.* 2008). Researchers tracking and modeling animal movement can extract behavioral states from telemetry and associated datasets, test hypotheses about what best predicts animal behavior, and explain how individual behavior scales up to landscape-level patterns of animal distributions. Recent advances in telemetry technology (Kays *et al.* 2015), the development of corresponding analytical methods (Long & Nelson 2013), and the integration of complimentary datasets (e.g., acceleration data; Wilmers *et al.* 2015; Spiegel *et al.* 2015a) have all dramatically increased movement ecologists’ inferential power. Especially in light of these developments, ecologists can decompose the impact of individual behavioral heterogeneity on pathogen spread with much greater ease, making movement ecology a promising avenue for exploring the behavioral underpinnings of how and why diseases spread in wildlife.

Both movement and disease originate in animal behavior at the individual level, and a feedback loop between the two emerges over time at broader ecological scales. For example, ecological theory suggests that the source-sink dynamics that naturally emerge between high-and low-quality habitat (respectively) can be reversed by an environmentally-transmitted disease, which turns high-quality habitat into an ecological “trap” (Leach *et al.* 2016). In practice, animal movement is driven by decisions that balance this trade-off between habitat quality and disease risk, and behavioral polymorphisms might even evolve as a consequence (Getz *et al.* 2015). For example, in an anthrax-endemic region of Namibia, zebra (*Equus quagga*) demonstrate a pattern of partial migration, where dominant herds appear to migrate away from high-quality habitat during the anthrax season, leaving behind lower-ranking resident herds to graze despite the higher disease risk (Zidon *et al.* 2017). Researchers posing questions solely about movement (why would zebra migrate away from high quality habitat?) or disease (why do some zebra select for areas with higher anthrax exposure risk?) would miss the overall pattern.

Understanding ecological links between movement and disease has direct implications for the way researchers model, forecast, and simulate wildlife disease outbreaks. The most basic models in epidemiology treat disease transmission as a function of the number of healthy and infected individuals in a population, linked by a transmission parameter (*β*). Doing so implicitly combines contact rates and transmission efficiency into one rate (McCallum *et al.* 2017), but individual heterogeneity in both is universally recognized as an important contributor to disease dynamics in humans (Lloyd-Smith *et al.* 2005a) and animals (Paull *et al.* 2012), and heterogeneity in movement can be an important predictor of this variation (Spiegel *et al.* 2017a). Where tools in movement ecology can help measure, describe, and predict heterogeneity in transmission between hosts, there are opportunities to pose novel questions relating to the effects of movement on contact (e.g., how do social networks structure contact rates?), the effects of contact on transmission (e.g., how does duration and proximity of contact affect the pathogen dose transmitted?), and the impact of infection on movement (e.g., does infection decrease or increase future contacts?). According to appropriate complexity methods in modeling (Larsen *et al.* 2016; Getz *et al.* 2017), the degree to which movement data should be incorporated into disease models depends on the kinds of questions being asked; but simultaneously, the resolution of available data on both movement and disease, and the level of prior knowledge, constrain the questions that ecologists can feasibly answer (Figure 1).

**Figure 1:**
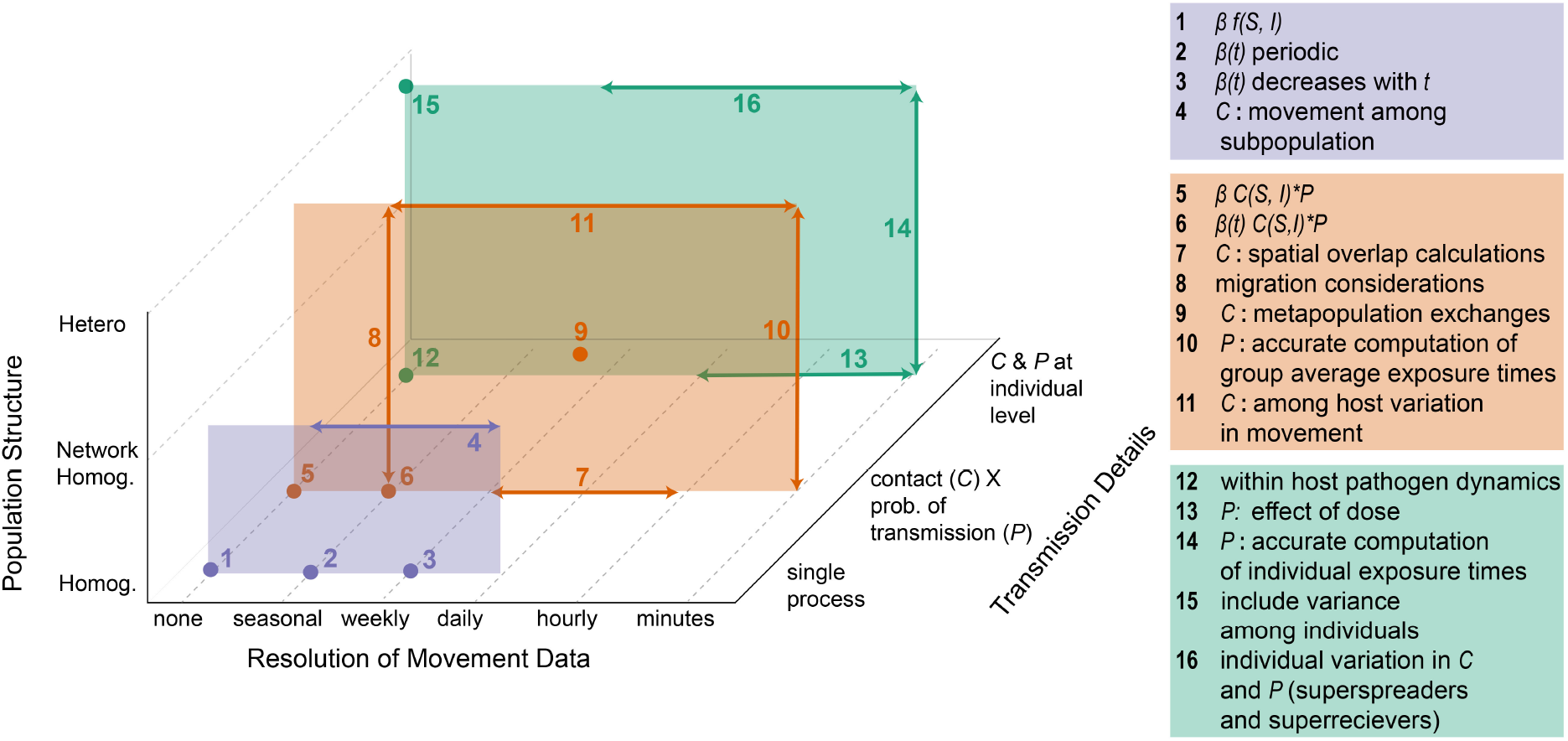
A Movement-Focused Modeling Space in Epidemiology. Incorporation of movement at different temporal scales can be used to address pathogen transmission-related questions at three levels (vertical colored planes). Transmission can be treated either as a single process (as commonly done in SIR models, where S are susceptible, I infectious and R removed individuals—see Box 1; purple plane), a concatenation of a contact process C and probability P of pathogen transmission during contact averaged over individuals (orange plane), or implemented at an individual level (green plane). In addition, transmission can be considered to occur within a homogeneous population, a network of homogeneous groups or subpopulations (metapopulation), or a spatially continuous heterogeneous population. Each labeled dot indicates a unique level of complexity that can be incorporated into the transmission process, while the spanning arrows imply that additional complexity can be incorporated at several different temporal scales (horizontal arrows) and population-structures (vertical arrows).

Here we synthesize the main ways that movement data are currently used to shed light on the processes underlying disease transmission, connecting animal behavior to broad patterns of wildlife (and human) health. Researchers unfamiliar with one or both fields are encouraged to refer to Boxes 1 and 2 for short primers on disease and movement ecology, respectively. We begin by describing how tools and methods from movement ecology can inform our understanding of how movement affects disease, potentially improving epidemiological models by better representing behavioral variation. Subsequently, we explore a more tentative application showing how movement data might directly improve disease surveillance. Throughout, we emphasize case studies that have successfully applied movement-based methods in these ways, and comment on particularly unexplored avenues and underutilized tools. Finally, we highlight the current state of synthesis work at the intersection of movement and disease ecology, and discuss the advances in data and models needed to move the field forward. In doing so, we recommend relevant movement ecology tools for studying processes underlying disease transmission (Table 1), and conclude by highlighting the broader implications for conservation and human health.

**Table 1:** Connecting Movement Methods to Disease Outcomes.

| Method | Description | Input data | Methodology | Benefits and insights | Reference |
| --- | --- | --- | --- | --- | --- |
| Spatial overlap | Estimate spatial overlap among hosts from their shared space use (a.k.a. passive interactions) | Coarse-grained spatial locations from multiple individuals | Various home range indices: minimum convex polygon, kernel density estimators, and local convex hulls | Identifying zones (or spatial scales) of potential influence and evaluating the likelihood of transmission for (primarily) indirectly transmitted pathogens | Yockney *et al.* 2013; Farnsworth *et al.* 2006 |
| Habitat selection | Environmental covariates can be used to predict areas of increased exposure risk based on the correspondence of host movements with models of vector/pathogen habitat preference | Host resource selection data; environmental predictors (e.g., landcover, temperature, water availability); vector/pathogen habitat suitability data | Resource selection functions are used determine habitat preference of host and parasite | A spatially explicit map of risk of exposure and transmission | Ragg & Moller 2000; Morris *et al.* 2016 |
| Behavioral analysis | Movement data can be used to identify specific behavioral states or canonical activity modes that affect the risk of exposure or transmission. Overlaying these behaviors on maps of environmental heterogeneity in pathogen prevalence can provide behavior-dependent risk maps | High resolution movement data or complementary behavioral data (e.g., accelerometer); maps of pathogen prevalence | Various methods to identify behavioral states from movement (and associated) data | A spatially-explicit prediction of the effects of behavior on infection risk (i.e., behavior- or individual-specific risk maps) | Gurarie *et al.* 2016; Nathan *et al.* 2012 |
| Network analysis | Calculate co-occurrence of individuals, dyads (a.k.a. dynamic interactions), and members of fission-fusion groups as a proxy for transmission | Spatial proximity or simultaneous spatial tracking of multiple sympatric individuals | Proximity sensors or fine-scale movement data to build networks from which various metrics can be derived | Identify direct contacts in the population as a proxy for transmission; isolate variation among individuals | Cross *et al.* 2005; Hamede *et al.* 2009; Silk *et al.* 2017a |

#### Box 1. A Disease Ecology Primer

Disease ecology as a discipline is conventionally focused on understanding the ecological drivers o *epidemiological* dynamics, referring to the study of the occurrence, distribution, and control of disease Whereas epidemiology conventionally focuses on human disease (including non-infectious causes o morbidity and mortality), wildlife epidemiology, and more broadly disease ecology, take a system; perspective on drivers of *infectious diseases*, those which are contagious within a population. Infectious diseases are spread by a *pathogen*, perhaps the most generic term for a bacterium, virus, or othei infectious agent (microorganism or *prion*) that can cause disease. Pathogens also include *parasites* a term defined ecologically that includes organisms that live in (*endoparasites*) or on (*ectoparasites*) another organism—its *host*—and benefit by deriving nutrients at the host’s expense. Not all parasites are immediately pathogenic (i.e., disease-causing). Some, such as ticks, could instead be the *vectors* thaï spread infectious agents, such as the bacterium that causes Lyme disease. Pathogens and parasites ar< spread by some process of *shedding*, the release of pathogenic material from a host either through passive emission (e.g., HIV in semen) or actively-induced emission when the life cycle of a parasite requires its own ejection from the host (e.g., aerosolization through coughing and sneezing or the fecal release o tapeworm eggs from a host). Some hosts, termed *super-spreaders*, can be particularly active shedders and infect disproportionately more susceptible individuals than other hosts do. In cases where shedding reaches a new host and this *exposure* event leads to infection, this produces an *effective contact*; what is considered an effective contact will vary with the mode of transmission of the pathogen in question.

In both humans and wildlife, outbreak dynamics are most readily modeled using a mathematical *compart-mental systems* framework: after dividing the population into epidemiologically-relevant compartments (*viz.*, susceptible: S, infected: I, and recovered: R), difference or ordinary differential equations are usee to describe the transitions of individuals between the disease classes over time. Typically these models make an assumption of *spatial homogeneity*, *random contact* among individuals, and *rapid mixing* of individuals within compartments. The course of infection is typically summarized for populations either via an *incidence* (the rate at which new cases arise), or *prevalence* (the proportion of the population infected) curve. If at least a low level of prevalence is maintained at all times, a disease is considered *endemic.* Ir contrast, an *epidemic* starts from a handful of introduced or new *index cases* and spreads throughout a susceptible population as an *outbreak*, before burning itself out. The latter occurs because the proportion of susceptible individuals in the population has either dropped below a *threshold density* or individuals have altered their behavior to avoid contact with infected individuals. When an epidemic is truly globa (defined by infection across multiple continents), it is referred to as a *pandemic.* In wildlife, *epizootic.* and *enzootic* serve as parallel terms to epidemic and endemic. Diseases that originate in wildlife and spread to humans are termed *zoonoses*, and are conventionally of special interest in disease ecology. The process of *spillover* of zoonotic disease into human populations is complex, and often poorly understood due to the complexities of human-wildlife contact. Conversely, *spillback* refers to the process by which a zoonotic disease is introduced by humans into novel animal host populations (whether domesticated or wild).

#### Box 2. A Movement Ecology Primer

Movement ecology has developed as a field that draws on *telemetry data* to explore the causes mechanisms, and patterns of animal movement, as well as understand its consequences on the ecology and evolution of individuals, populations, and communities. *Telemetry* refers to the process o transmitting and recording the positions of an animal, and represents the primary means of detecting animal movements. Early telemetry research relied upon *Very High Frequency* (VHF) radio signal; to triangulate the positions of *collared* or *radiotagged* individuals. The individual positional fixes obtained can be referred to as *relocations*, and when treated consecutively, they are often called a *trajectory* or *path.* The relatively coarse temporal resolution of most relocation data from classica radiotagging methods limits the ability of movement ecologists to observe and differentiate among *fundamental movement elements* (FMEs; e.g., a walking versus trotting step) that make up the movement path of an individual. However, the relatively infrequent or irregular fixes emerging from such devices can still be used to evaluate patterns of *space use* and *habitat selection*. For example even coarse movement data can aid in characterizing the manner in which an animal utilizes its *homi range*, which represents the area it traverses in its daily activities of foraging, mating, and caring for young. These areas have been delimited in a number of ways, including: *minimum convex polygon* (MCP) methods, which simply construct a boundary around the outermost points of a trajectory; anc *utilization distribution* (UD) methods, which offer more information regarding the frequency of space use within the home range. Recently, alternative methods that more explicitly account for the tempora component of movement data have been proposed, including the *time-local convex hull* (T-LoCoH) method and *Brownian bridges*, among several others. Even with sparse datasets, these methods are expected to create meaningful generalizations of space use and can form the basis of *spatial overlap* analyses that aim to determine the level of shared space use among monitored individuals. Severa methods for understanding individual and population level habitat selection, such as *resource-selection functions* (RSFs), can also be used with relatively coarse movement data. These methods aim to identify the habitat types that an animal prefers, indicated by disproportionately greater use of a habitat than expected based on its availability on the landscape, and create predictive maps of space use

Today, the majority of movement ecology research depends upon more advanced satellite technology referred to broadly as *Global Positioning Systems* (GPS), to record animal locations at finer spatial am temporal resolutions. Even with this technology, consecutive relocations typically span a mix of FMEs Nonetheless, a variety of summary metrics can be used to describe the path, the most basic of which are the *step length* (the Euclidean distance between consecutive relocations) and *turning angle* (the angle of one step relative to the step immediately prior). The higher resolution relocations can also enable *behavioral analyses*, which often rely on *path segmentation* methods to split a movement trajectory into segments that look quantitatively similar (often based on those simple summary metrics). Such analyses can help determine the *behavioral state* of an individual at specific points in time. These states occur at coarser time scales than FMEs, but represent short-lived phenomena that can be inferred from GPS data Similar analyses can allow for the clustering of longer sequences of behavioral states that are considered collectively as *canonical activity modes* (CAMs; e.g., resting or foraging), which are also readily observable in modern telemetry data. For example, foraging is a CAM that often consists of a variety of behaviors including searching, eating, and perhaps vigilance, among others. A full movement path, however often consists of a series of CAMs, and *movement syndromes* are used to describe movement patterns at the scale of an entire trajectory, enabling discrimination among types of individuals (e.g., territoria versus nomadic individuals). With recent technological advances to the telemetry units worn by animals supplementary data sets, such as those obtained using *accelerometers* that measure changes in velocity in three-dimensions, have enabled the evaluation of movement behaviors at even finer spatiotempora scales, getting researchers closer to observing FMEs. Similarly, the advent of *proximity sensors*, whicl record when two collared animals are within a specified distance of one another, has allowed researchers additional insight into the *spatial proximity* of monitored individuals. These data can be used to inform *contact networks*, which map the associations among individuals in a population.

## Movement affects Disease

Depending on a pathogen’s mode of transmission, different tools in movement ecology will be more or less suitable for exploring transmission risk. We make the broadest possible division, placing pathogen life histories along a spectrum between direct and indirect transmission (Figure 2). Direct transmission refers to pathogens that require contact between an infected and susceptible animal at the same place *and* at the same time. Indirect transmission, on the other hand, describes pathogens that can occupy some intermediate reservoir or vector between hosts (i.e., a host of another species, or an environmental reservoir like soil or water), making spatial overlap a more significant requirement than temporal overlap. Whether a pathogen is treated as directly or indirectly transmitted should depend on both the duration of time it can survive outside of hosts, and its ability to disperse in the environment separate from host movement. Temporal overlap between animals matters less when infective stages survive for extended periods outside of hosts, or when the infective stage moves independently (e.g., when environmental forces induce relatively long-distance dispersion, a feature common in marine systems where pathogens are often at the mercy of currents; Lafferty 2017).

**Figure 2:**
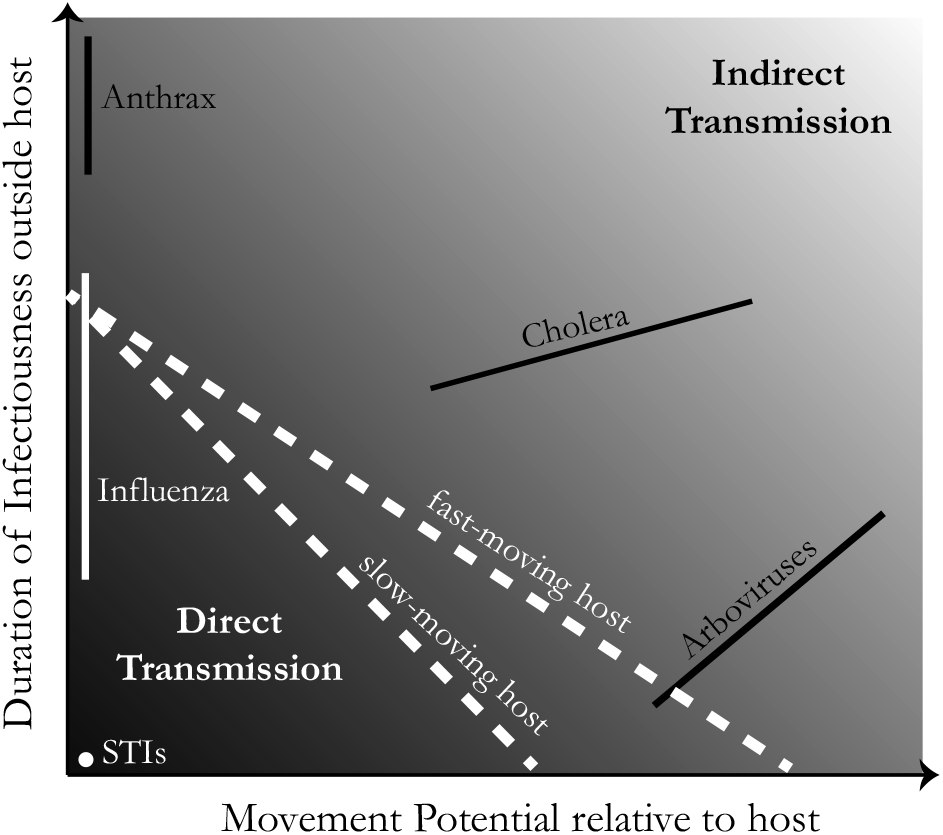
The Transmission Continuum. Transmission mechanisms vary across a continuous spectrum. The classification of a particular pathogen or parasite in a given system depends on the movement potential of pathogens relative to their hosts and the ability of pathogens to remain infectious outside hosts. Those pathogens that require two agents to interact directly for successful transmission, often via a specific behavior, such as sexually transmitted infections (STIs), are an unambiguous example of a directly transmitted disease and represented by a point. Pathogens that transmit successfully over a broader set of conditions, such as influenza or arboviruses, are represented conceptually across the gradient as a line and might vary across one or or both of the axes. Along this spectrum, we have determined a somewhat subjective threshold between what we describe as *Direct transmission* and *Indirect transmission*, visualized by the white dashed lines. Even within the same pathogen taxon (and thus, the same characteristic duration of infectiousness), this threshold could shift along this gradient depending on the relative speed of host movement.

Broad categories of infectious agents (bacteria, viruses, parasites, etc.) are unlikely to map neatly onto direct or indirect transmission. For example, some ectoparasites are directly transmitted among members of a social group (e.g., some species of avian lice; Rózsa *et al.* 1996), whereas others often spend time freely moving off-host (e.g., several tick species that infect reptiles; Sih *et al.* 2017). Some pathogens may also alternate between direct and indirect modes; for example, Zika virus and canine leishmanisis are both vector-borne diseases with rare sexual transmission events. Similarly, influenza is usually directly transmitted through air or direct contact, but can sometimes persist in the environment via fomites (nonliving object or substance capable of carrying infectious material) for hours or days (Weber & Stilianakis 2008). Whether researchers choose to focus on spatial or spatiotemporal overlap, corresponding to direct or indirect contact, is likely to depend on the scale at which other host processes are modeled, and the spatial and temporal extent of the analysis (see Box 3).

#### Box 3. Balancing Scales: Simultaneous Modeling of Movement and Epidemiological Processes

Deciding on a spatiotemporal scale for epidemiological models is usually a function of the timescales of host and pathogen processes, including temporal aspects of transmission like latency, persistence in the environment, or replication rates during early infection. The rates of biological processes might not map directly onto the models we build, if the temporal scale of data is necessarily coarser due to the resolution of available movement data. However, some of the greatest successes of movement ecology have involved explicit model formulation with attention to spatiotemporal resolution of processes (Lyons *et al.* 2013) potentially offering a template for integrated work. We outline some brief guidelines:

**Space pixel.** The corresponding spatial resolution *S* is related to time through the diffusion relationship:

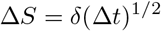 This is where movement comes in: *δ* is a movement diffusion constant estimated from empirical data and will vary among organism types. An alternative approach is to use a velocity relationship

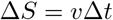

for organisms that mainly execute directed movement at average velocity v at fine time scales Since empirical tracking data has repeatedly shown that movements of animals (and humans) are often super diffuse, we suggest that former approach as generally more favorable (Raichlen et *al* 2014; Spiegel *et al.* 2015a).
**Coarse graining.** Going from the scale of individual transmission upwards to emergent processes like landscape structure, epidemic wavefronts, or even range shifts requires proportionally aggregating data and model structure, a process typically termed *coarse graining.* There are various levels of coarse graining, each representing close to a one magnitude of size step up. Coarse graining require; aggregating over a union of pixels, using an appropriate integral kernel. Integral kernels can take several forms including the bounded uniform, truncated Gaussian, or other more idiosyncratic choices. Optimal kernel choice can be guided by wavelet analysis of movement data.
**Time pixel.** The minimum time resolution should be based on some fraction of the most fundamental cycle pertaining to the problem; for example, movement data might be recorded at a resolution Δ*t* of every 15 minutes, though this is much shorter than the typical interval in epidemiological models.
**Temporal scaling.** Increasing scales of temporal aggregation can provide different results and absorb more noise in data by matching biologically-relevant timescales. For example, if Δt is 15 minutes then 100Δt is approximately one diurnal cycle, 3,000Δt is approximately a lunar cycle, and 10,000Δt is approximately the length of one season in a four season year. Models can also b< downscaled, which might be appropriate under highly data intensive conditions. However, as finescale processes emerge at finer scales, models might lose predictive accuracy without incorporating finer data or processes.
**Appropriate complexity.** At various spatiotemporal levels of resolution different epidemic models might apply and the question arises as to the appropriate level of complexity in the model (Larsen *et al.* 2016). For example, models should include a within-host component at the level of Δt =15min, while epidemiological models might include daily rates of detection and isolation of individuals at diurnal levels of resolution, or could include transmission rates that exhibit seasonal variation if epidemics last several months or more. Additionally, incorporation of movement into models might require individual-based approaches for the finest scales of analysis (Getz 2013).
**Multiscale modeling.** As data and models are aggregated, models can be run to reflect the multiple timescales on which movement and epidemiological processes operate. Wavelet decomposition of movement data can inform the most important concurrent scales of movement processes; similar analyses can be performed with time-series epidemiological data, when available.

## Direct Transmission

Directly transmitted pathogens rely on contact between infected and susceptible individuals. Contact rates (process *C* in Figure 1) are most easily thought of based on the frequency and strength of interactions between animals in a population, a problem that lends itself naturally to network methods (Silk *et al.* 2017a,b). Meanwhile, the probability of transmission during contact (*P* in Figure 1) will depend largely on the duration and nature (e.g., grooming vs. fighting) of the contact needed for pathogens to spread, which can be incorporated into network analyses in various ways.

Networks are a statistical model that abstract population structure as a set of connected nodes, traditionally representing individual animals in the population. Edges indicate the connections between individuals, whether these are defined as interactions of a certain duration, or individuals coming within a certain distance of one another. Such information can be displayed graphically through the use of directionality (arrows) or weight (line thickness). Directionality could indicate an epidemiologically-relevant behavior that impacts the actors differently (e.g., grooming), while weight can be derived from the frequency or duration of such interactions (Cross *et al.* 2005). The components of a social network may ultimately be spatially implicit (i.e., animals’ position in the network cannot be projected onto a map), but these networks can be informed by movement data in cases where in-person behavioral observation is impractical or infeasible, making them a valuable tool for reconstructing the spread of directly-transmitted disease. Networks can also be constructed in the context of indirect transmission, but might require different data (e.g., capture histories from an array of traps; Davis *et al.* 2015) or the inclusion of a time lag to emphasize the spatial component of transmission (e.g., Sih *et al.* 2017). For a visual example of these concepts, see Figure 3.

**Figure 3:**
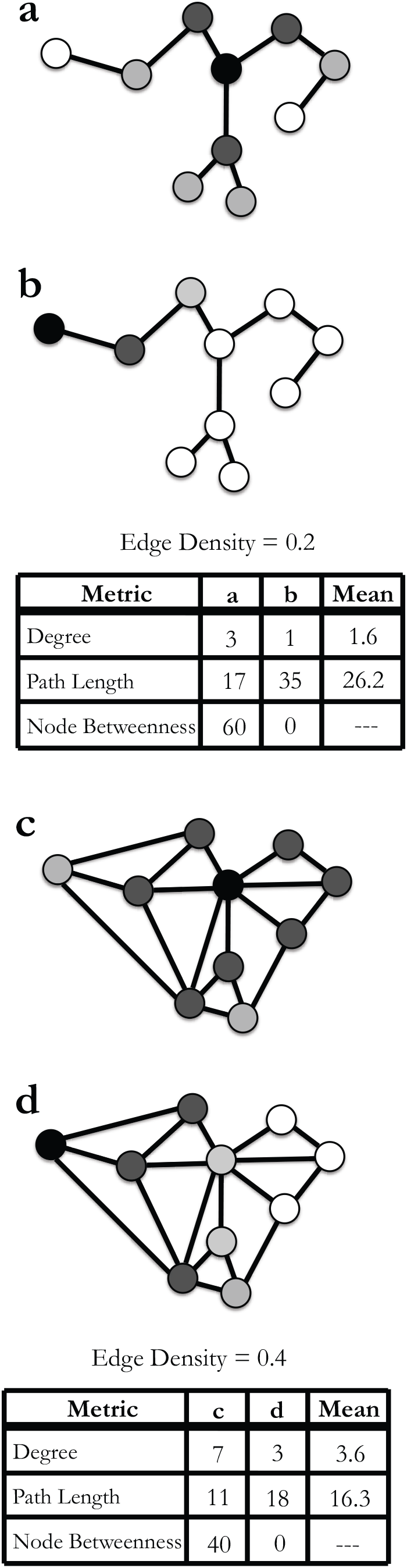
Networks for Disease Research. Network analyses can serve to identify particular contact network structures that might be conducive to disease spread through a population and identify individuals within networks that make disproportionate contributions to the transmission of a disease (Ryan *et al.* 2013). A network with relatively low edge density and high path lengths might prevent a directly transmitted parasite (or pathogen) from spreading through a population (networks **a** and **b**). Contrastingly, a network with high edge density and low path length could facilitate parasite spread through a population (networks **c** and **d**). In addition, the position of the first infected individual (shaded in black) in a network might facilitate or inhibit a parasite from spreading. Individuals with relatively high degree or node betweenness could be super-spreaders (networks **a** and **c**), whereas individuals positioned at the periphery of a network, with lower degree and node betweenness, might cause transmission to fade out (networks **b** and **d**). At both the population and individual levels, these network characteristics depend on resource distribution, social relationships, and ultimately, the movement behaviors that arise from both. It should also be noted that the same general principles would apply if this schematic were imagined as a spatial network instead of a contact network, with nodes representing locations rather than individuals.

Most networks extracted from movement data are proximity based social networks (PBSNs). They can be constructed using either special proximity sensors, or from movement data using a spatiotemporal threshold value to designate contact between animals (e.g., within *M_c_* meters for at least *T_c_* time units; Farine & Whitehead 2015). Observed association patterns in social networks are often compared to expected patterns in null models (e.g., ideal gas model) or randomized networks, to test hypotheses about the mechanisms underlying social structure (Farine 2017; Silk *et al.* 2017b). For example, by randomizing the order of daily movement paths *within* each individual, rather than *between* individuals (as is typical in most network randomization methods), Spiegel *et al.* (2016) developed a method to assess sociality separate from associations resulting from the spatial structure of the environment. An extension of this approach allowed for the identification of the locations of interactions and revealed the sex-specific patterns underlying the network structure (Spiegel *et al.* 2017b). These networks have been a key part of efforts to understand how ticks are transmitted in sleepy lizards (*Tiliqua rugosa*), reptiles with an unusual life-long pair breeding pattern that may facilitate tick transmission (Sih *et al.* 2017).

Social networks can provide insights into disease spread even in the absence of explicit disease data (Craft & Caillaud 2011). Different species’ social behavior may correspond broadly to different network structures, and corresponding outbreak dynamics; for example, social hierarchies may comparatively limit the rapid spread of epidemics, whereas “gregarious” species with connected, unfragmented social networks are prone to major outbreaks (Sah *et al.* 2017b). At the population level, the overall characteristics of a network (e.g., average degree of nodes, path lengths, and edge densities) can be vital for understanding the hypothetical implications for transmission (Craft 2015), including vulnerability to epidemic spread (Porphyre *et al.* 2008; Craft *et al.* 2011). In a meta-analysis, Sah *et al.* (2017a) found that modularity (i.e., the strength of division of a network into separable components) has a surprisingly limited effect on outbreak size and duration, especially for higher levels of modularity. However, fragmented networks with high subgroup cohesion still experience comparatively limited and brief outbreaks. In a relevant case study, Hamede *et al.* (2009) used proximity sensors to build a comprehensive contact network of Tasmanian devils (*Sarcophilus harrisii*) in a population at risk from the introduction of a directly-transmitted parasitic cancer. The entire population was connected in a single network, allowing the spread of a pathogen from a single individual—and therefore, preventing most containment efforts in the event of an outbreak (Figure 3).

At the individual scale, networks can show where individual heterogeneity in transmission occurs (Lloyd-Smith *et al.* 2005b; Perkins *et al.* 2009; Paull *et al.* 2012). Similar metrics to those employed at the population level can also describe single nodes or edges within a network, potentially illuminating differences among individuals within a population (White *et al.* 2017; Silk *et al.* 2017a). For instance, Weber *et al.* (2013) found that degree (the number of connections a given node has to other nodes), closeness (effective distance between an individual and all others in the network), and flow betweenness (a measure of the role of a particular node in connecting all other pairs of nodes in the network) were associated with tuberculosis infections in badgers (*Meles meles*). Because causality could not be determined, the researchers concluded that either an individual’s network position could affect infection risk, or that infection could affect network position. By showing how heterogeneity among hosts propagates an infection through a susceptible population, analyses such as these could help identify super-spreaders, which in turn could help improve estimates of *R_0_* (i.e., the expected number of secondary cases produced by a single infection in a completely susceptible population; see Box 1; Lloyd-Smith *et al.* 2005b).

The use of proximity data synchronized with GPS and accelerometer data can help better identify social interactions that are epidemiologically-relevant (Nathan *et al.* 2012; Brown *et al.* 2013). Some pathogens require sexual contact for transmission (like herpes viruses), whereas others need only a brief physical contact (like influenza). In this sense, movement-based behavioral analyses can decompose sociality into interactions with implications for disease transmission, improving the relevance of network analyses. Even without network data, movement analyses might identify behaviors that can be linked to interactions among individuals (Bartumeus *et al.* 2005; Fryxell *et al.* 2008) or to the social standing of individuals (Wittemyer *et al.* 2008), allowing for inferences about the vulnerability of individuals to disease. For example, Wittemyer *et al.* (2008) used wavelet analysis of three-hourly location data to infer that the social rank of elephants (*Loxodonta africana*) affects the periodicity of their movement at a multiday scale. In addition, they found that lower social standing correlated with higher movement variability during the resource-deficient dry season. This and similar analyses can be used to identify which individuals might interact most frequently (here, based on social rank). They could also be used to identify individuals whose irregular access to resources stresses them to the point where they become vulnerable to infection. Social structure could influence susceptibility in other ways (Altizer *et al.* 2003). For example, social rank can determine the form and frequency of breeding behaviors in the group, making it especially relevant for sexually-transmitted infections. Additionally, social living could confer anti-parasite benefits such as increased parasite resistance or tolerance (e.g., due to regular or low dose transmission between conspecifics), or could mitigate disease (e.g., due to increased fitness as a result of superior resource acquisition in a group; Ezenwa *et al.* 2016).

## Indirect Transmission

In the case of pathogens and parasites that are transmitted indirectly (Figure 2), the processes by which a one host sheds a pathogen and another host is exposed are independent and might rely upon different host behaviors (e.g., defecation for the former and foraging for the latter). Tools from movement ecology offer a way to consider these processes separately from the perspective of the infected individual and susceptible individual at various time scales (sub-hourly to multiweek time, as depicted in Figure 1).

High resolution movement data (i.e., sub-hourly: Figure 1) enable researchers to estimate the frequency and duration of encounters with known pathogen hotspots on a landscape (e.g., mosquito breeding sites at standing water). Though practical considerations might limit the number of animals that can be monitored in a study population (Williams *et al.* 2014), appropriate sampling schemes offer a basis for statistical inferences that apply more broadly. For example, existing tools can identify associations between habitats or time periods and animal presence, thereby offering insight into overlaps with infectious sites (Figure 4). Further, if movement data help identify behavioral drivers (e.g., resource distribution and its seasonal changes), then insights from the monitored subset of the population could be used to mechanistically model encounter probabilities or factors contributing to shared space use (e.g., Cross *et al.* 2005; Spiegel *et al.* 2015b).

**Figure 4:**
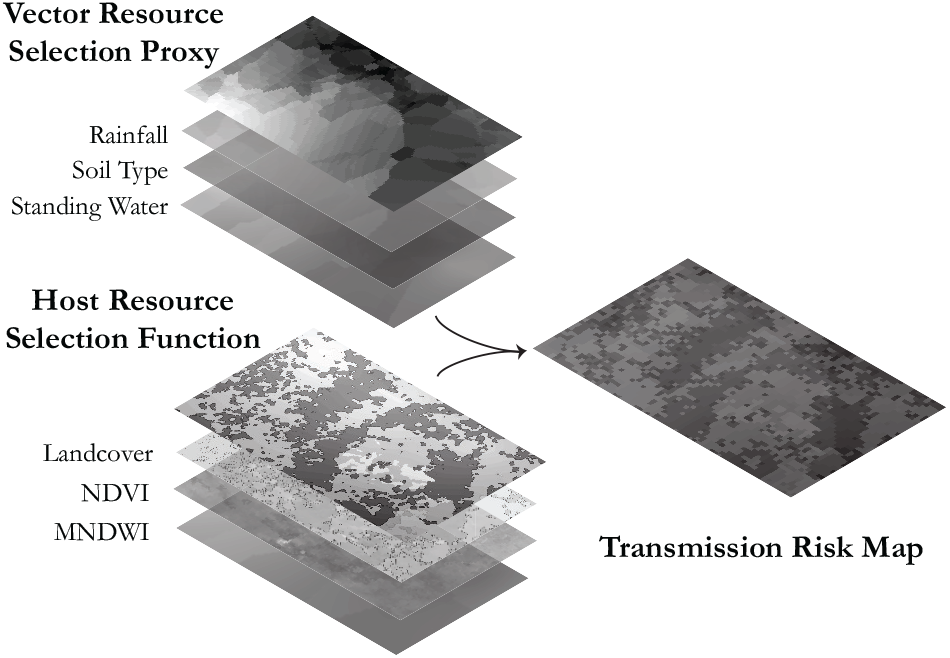
Calculating Spatial Risk from Movement Data. For vectors with known associations to abiotic covariates, resource selection functions can be a powerful tool to identify areas of overlap with host movement and map areas of increased exposure risk. In this hypothetical example of an arbovirus, maps of resource selection or association (the top layer of each stack) are derived for a terrestrial host and a water-dependent vector from associated environmental layers (e.g., land cover or soil type) and movement or presence data. Combined, these maps of resource selection can produce a map of overall transmission risk. Alternatively, a similar approach could be used with a pathogen, such as anthrax, that relies on mappable soil characteristics, such as calcium levels and pH (Mullins *et al.* 2013). The other layer would correspond with host habitat preferences, including indicators of watering hole locations (i.e., Mean Normalized Difference Water Index; MNDWI) and graze or browse quality (i.e., Normalized Difference Vegetation Index; NDVI).

Clustered observations reflect spatial regions that individuals frequent, and can indicate areas where encounters among individuals (tagged or untagged) are more likely. Applying techniques to identify such clusters in data from multiple animals (Webb *et al.* 2008; Seidel & Boyce 2015; Van Moorter *et al.* 2016) can aid in identifying population-wide aggregation points with potential epidemiological significance. These aggregation points might reflect underlying environmental heterogeneity (e.g., waterholes) or social contacts (e.g., leks) (McNaughton 1988; Carter et *al.* 2009); regardless of the mechanism driving aggregation, these locations are likely to be important for estimating relative exposure risk. Various methods can help distinguish social and environmental causes of such aggregation patterns (e.g., Spiegel *et al.* 2016; Borchering *et al.* 2017), potentially offering a way to assess transmission risk.

Areas of dense use are also identifiable through the construction of utilization distributions (UD), which illustrate the relative frequency distribution of the location of a particular individual over time (Van Winkle 1975). UDs are most commonly derived using kernel density estimation techniques (Worton 1989). Methods for estimating space use at broader scales, especially estimates of seasonal range size and overlap, have been included in epidemiological models. For example, Ragg & Moller (2000) used radiocollars, in conjunction with other methods, to track the microhabitat selection of both active and denning feral ferrets (*Mustela furo*), a vector of bovine tuberculosis (*Mycobacterium bovis*) in New Zealand. Ferret movements were found to be concentrated in grazed areas and at ecotones between pastures and vegetation cover, thereby increasing their risk of transmitting tuberculosis to possums and livestock. Similarly, Conner & Miller (2004) used cluster analysis on mule deer (*Odocoileus hemionus*) location data to identify population units, and used kernel density estimation to delineate seasonal ranges for each population. Subsequent analysis showed that winter ranges rarely overlapped (< 1%), likely due to their smaller size, whereas summer ranges had >22% overlap among population units. Therefore, researchers concluded that summer ranging behavior was likely responsible for the spread of CWD among subpopulations, whereas winter ranging behavior had the potential to amplify CWD prevalence within a subpopulation if an infected individual was present. In an extension of the study, Farnsworth *et al.* (2006), used area estimates of summer, winter, and individual home ranges to frame regression models at different scales. They found that movements within individual home ranges had the greatest implications for CWD exposure, highlighting the potential of high-resolution movement data to alter our understanding of the mechanisms underlying observed patterns of transmission.

Novel methods that consider the temporal autocorrelation inherent in movement data enable more detailed home-range delineations than those that emerge from traditional, purely spatial, estimators (Benhamou & Riotte-Lambert 2012; Lyons *et al.* 2013). Additionally, these methods might produce more accurate results when home-range overlap is used as a proxy for exposure risk, especially in cases where the pathogen’s ability to survive outside a host is limited. One such method, time-local convex hulls (T-LoCoH; Lyons *et al.* 2013), creates time-dependent hulls within the utilization distribution from which various metrics can be derived. Two such metrics are the duration of a visit to a particular point or area of interest, known as the residence time, and the rate at which individuals return to them, known as the visitation or return rate. Used together, these metrics can offer a means of evaluating the relative risk of contact or exposure among individuals (Dougherty *et al.* 2017). Site-fidelity metrics such as these could be particularly important in the case of indirectly transmitted pathogens, because high levels of fidelity increase exposure risk if an infectious reservoir is present in the range, but will buffer an individual from exposure if the range is free of relevant pathogens or parasites. Thus, higher mean visitation and duration rates should indicate greater heterogeneity of infection risk across individuals in a spatially-structured population.

Beyond general descriptions of space use, tools that explore landscape level patterns and probability of use—which are some of the most developed in movement ecology—can offer predictions regarding where susceptible individuals might be exposed to disease. Habitat-selection methods, such as resource-, path-, or step-selection functions (RSF, PSF, and SSF, respectively), can illuminate landscape features and types preferred by individual hosts or the population as a whole (Leclerc *et al.* 2016). These methods, used to infer the probability of use of any given resource unit within the range of a population, quantify which habitats animals select within their range (Boyce & McDonald 1999; Manly *et al.* 2002). By comparing points used by animals in the population to those available but unused within their range, RSFs provide a statistical model of habitat preference (Boyce *et al.* 2002). In the context of disease, these models can identify habitats where pathogen deposition and, thus, exposure are most likely to occur based upon their relative probability of selection. For example, Morris *et al.* (2016) built an RSF for elk (*Cervus elaphus*) ranging in the presence of soil-borne anthrax (*Bacillus anthracis*) in southwestern Montana. Based on the preferences of the elk and a parallel evaluation of the landscape features that enabled long-term persistence of anthrax spores (with ecological niche modeling), Morris *et al.* (2016) mapped the areas of highest risk to the elk population.

In cases where pathogens or parasites are difficult to study but follow predictable patterns of occurrence on a landscape, RSFs and other movement tools could allow researchers to identify potential hotspots for vector-borne or environmental transmission (Figure 4) using GIS technology. The application of GIS is particularly suitable when vector preferences on a landscape are well understood, as in studies of the use of fragmented forests near agricultural land by ticks (a vector for Lyme disease; Allan *et al.* 2003; Brownstein *et al.* 2005) or mosquito use of standing water for breeding sites (Perkins *et al.* 2013). The relevance of these approaches will be strongly dependent on how far vectors can move, as well as the importance of dispersal in the life cycle of vectors and the overall prevalence of disease. A similar application can easily be imagined for pathogens maintained in soil, such as anthrax (*Bacillus anthracis*) or plague ( *Yersinia pestis*); or in water, such as cholera (*Vibrio cholerae*) or cryptosporidiosis (*Cryptosporidium parvum*). The pathogens in all four of these examples follow predictable patterns of occurrence and persistence based on abiotic environmental variables (Carlson *et al.* 2017). The dual RSF framework helps researchers to identify whether host populations select for areas with high infection risk. In addition, such methods can indicate whether certain individuals are using these features more than others, offering insight into the heterogeneity of exposure throughout the population.

## Disease affects Movement

Movement tools may also provide a more direct (but underexplored) tool for disease surveillance, as infection often affects host behavior in observable ways. Pathogens can alter host movements either through vigor loss (i.e., the appropriation of resources towards an immune response) or host manipulation (direct chemical or physical modification by the pathogen). Examples of infection-induced behavioral shifts range from *Cordyceps* fungi in arthropods, which cause hosts to climb to the upper part of a plant before death (Roy *et al.* 2006), to *Toxoplasma gondii* in rats (*Rattus norvegicus*), which results in higher activity levels and loss of fear in infected hosts (Berdoy *et al.* 2000). Importantly, such changes can alter movement trajectories (Murray *et al.* 2015; Cross *et al.* 2016) in ways detectable by movement tools (e.g., risk-taking behavior or a dramatic shift in habitat preference), potentially allowing researchers to identify shifts in individuals’ behavioral patterns once individuals become infected.

Movement trajectories can be characterized by sets of metrics extracted from consecutive relocations. These include step length (the distance between two consecutive points), relative turning angle (the angle between the trajectory indicated by two points relative to that inferred from the previous step), and persistence (the tendency of a movement to persist in a particular direction). Since these telemetry data are discrete, if they are not sufficiently fine-scaled, they cannot be used to characterize fundamental movement elements (FMEs, Box 2; Getz & Saltz 2008). They can, however, be used to cluster movement path segments into canonical activity modes (CAMs; Figure 5; Getz & Saltz 2008) using thresholds, clustering, and behavioral changepoint techniques (Gutenkunst *et al.* 2007; Van Moorter *et al.* 2010; Gurarie *et al.* 2009, 2016).

**Figure 5:**
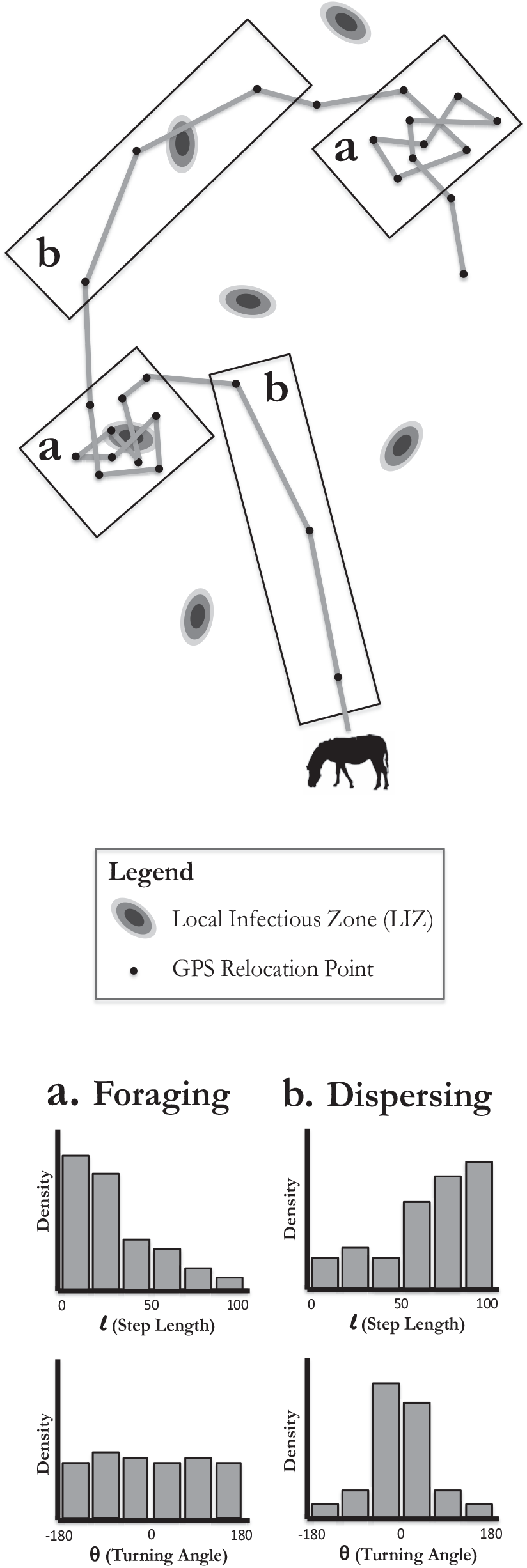
Canonical Activity Modes (CAMs) from Movement Data. Several alternative methods enable a researcher to infer different canonical activity modes (CAMs; thematic mixes of behavioral states). In this schematic, a hypothetical trajectory of a zebra can be easily divided into a foraging CAM (boxes labeled **a**), defined by relatively small step lengths and an almost uniform distribution of turning angles, and a dispersing CAM (boxes labeled **b**), defined by relatively larger step lengths and a distribution of turning angles with a low variance. For disease research, if a pathogen is known to have environmental reservoirs with predictable locations (e.g., due to its dependence on certain soil types or pH), the CAM during which the animal is susceptible (in this case, foraging, when the zebra eats plants or soil harboring the bacterium) can be isolated to identify the areas or times of greatest risk. One can also identify individuals or classes (e.g., sex or age groups) who could be at greater risk than others due to the higher proportion of time they spend foraging in their activity budgets. In this specific example, the host is at low risk of transmission from the LIZ in box **b** and at high risk from the LIZ in **a** due to the different behavioral states. The gray lines between GPS relocation points represent estimated paths between known locations rather than an exact trajectory.

The above movement trajectory metrics might differ sufficiently between healthy and infected individuals to allow them to be used to identify an individual’s disease state. Further, infection with a pathogen could affect daily activity budgets, potentially altering the number or distribution of change points seen across a day. The segmentation of movement paths into CAMs or, at a finer scale, behavioral states (Nathan *et al.* 2012), represents an active area of study in disease ecology (Edelhoff *et al.* 2016). For example, Cross *et al.* (2016) established that infection with mange (*Sarcoptic scabiei*) in wolves (*Canis lupus*) was associated with decreased daily movements, with later stages of infection reducing total distance more than earlier stages. In addition, infected wolves spent significantly less time in an active behavioral mode (defined as hourly movements greater than 50 meters) than healthy wolves, with degree of infection once again affecting activity level. Similar comparisons can be performed with data collected at a coarser scale, as exemplified by Murray *et al.* (2015), who demonstrated that disease state was related to differences in home-range size of coyotes (*Canis latrans*) infected with mange. Movement data derived from complementary sensors, on the other hand, offers researchers even deeper insight into the impacts of disease on movement behavior. Accelerometers, for example, enable the detection of tremors in individual paths and can help differentiate between bold versus submissive walking gaits, which can be indicative of different disease states. In a study of cockroaches (*Blaberus craniifer*), Wilson *et al.* 2014 extracted the vectorial dynamic acceleration (VDA; Shepard *et al.* 2008), a metric for characterizing the tremors in an animal’s movement, and found that the dynamism in each stride decreased with progressing fungal infection.

While the the application of movement ecology to disease diagnostics remains relatively unexplored, an ability to identify infected individuals from movement tracks could be highly useful in systems where diagnosis is difficult, invasive, or lethal (especially important for species of conservation concern). These methods might also enable researchers to infer the approximate onset time of symptoms, in turn improving disease models. The increasing availability of detailed movement data provides researchers an opportunity to develop and validate new methods along these lines.

## Synthesizing Movement and Disease

Ecology, as a scientific discipline, advances through the interplay of data, models, and theory: work at the interface of movement and disease ecology is rapidly growing on all three fronts. We briefly comment on how models can bridge data-driven understanding into theoretical results, and then present a systematic literature review showing the biases in how different movement tools are currently used to explain and predict disease dynamics.

## Scaling Models to Theory

Compartmental models (Box 1) are a nearly universal tool for studying human and wildlife diseases (Anderson *et al.* 1992; Keeling & Rohani 2008), and have been applied to a broad range of host-pathogen systems, with numerous extensions for host-age effects, pathogen-strain effects, or even the influence of pathogens on host behavior. Compartmental models, however, are not easily adapted to account for the effects of landscape and population spatial structures on *risk of infection* (Figure 4). Accounting for this level of variation requires a representative sample of individuals within the population to be tracked and their contact rates with other individuals (direct transmission) or infectious environmental locations (indirect transmission) recorded. Mechanistic models allow researchers to upscale individual patterns (such as behavioral rules or contact patterns) to a broader population, and are frequently used to validate or test experimental results. For example, disease outbreaks are easy to project on simulated networks, allowing researchers to confirm hypotheses about how modularity and fragmentation link animal social structure to outbreak size (Sah *et al.* 2017a,b). However, directly upscaling animal behavioral rules into spatiotemporal patterns of disease may require researchers to build individual-or agent-based models (IBM, ABM; Grimm *et al.* 2005).

More specifically, IBMs can use step length, turning angle, canonical activity mode distributions, habitat or resource preferences, or even various network-based metrics to generate likely movement paths for all individuals in the population. With basic assumptions about transmission rates as a function of contact duration, these trajectories can be used to simulate disease outbreaks on real landscapes with “real” animal movement principles. An number of IBMs that incorporate mechanistic movement rules to explore disease dynamics have been constructed (Bonnell *et al.* 2010; Dion *et al.* 2011; Tracey *et al.* 2014; Belsare & Gompper 2015). One of these (Bonnell *et al.* 2010) used individual host energy levels to generate movements toward higher resource patches. These foraging decisions ultimately drove microparasite transmission dynamics among red colobus monkeys (*Procolobus badius*) as they shifted their distributions on the landscape in search of food.

An obvious drawback of IBMs compared to compartmental models is the high computational demand associated with running simulations at this scale, though this limitation is becoming less prohibitive with the increasing availability of high performance computing. Perhaps a more serious limitation, IBMs involve many more parameters than compartmental models, thereby increasing difficulties associated with verification and validation procedures (Filatova *et al.* 2013). In addition, IBMs generally include stochastic elements, which can make statistical inference using IBMs very challenging (Hartig *et al.* 2011). While recent methodological advances have overcome some of these limitations, they remain impediments to the broader application of IBMs in disease modeling. Continued efforts to synthesize movement and disease ecology, however, are likely to inspire the development of new solutions for translating risk (based on movement behaviors on a specific landscape) into generally applicable rates for epidemiological models.

We also caution that mechanistic models (individual-based or otherwise) that explicitly incorporate movement rules from empirical data might not be transferable across space, or even across seasons or years. For instance, if environmental change alters behavior (e.g., annual migration targets shift in response to climate change), even mechanistic models based on empirical movement data might become inaccurate. This could be problematic for predicting pathogen dynamics in response to rare movement events (e.g., atypical long-distance dispersal events) or transmission (e.g., cross-species spillover events). Some tools exist in epidemiology to address model building based on limited data (e.g., fitting *R_0_* for rare spillover diseases; Blumberg & Lloyd-Smith 2013; Kucharski & Edmunds 2015), but this problem requires special attention in the context of movement research, and given the ongoing anthropogenic changes to local and global environments.

## Current State of the Synthesis

In a review of Web of Science, we found 70 papers published between 2000 - 2017 using movement tools in disease research (see Supplementary Appendix 1 for details). For the purposes of the review, we did not include agent-based modeling studies without empirical basis, though we noted they followed similar biases. This literature review revealed a notable bias across study organisms (Figure 6). Most studies focused on pathogens that can spillover to human and domestic animal populations, including bovine tuberculosis (*Mycobacterium tuberculosis*), anthrax (*Bacillus anthracis*), brucellosis (*Brucella abortus*), foot and mouth disease (FMD; *Aphthae epizooticae*), and chronic wasting disease (CWD). Hosts with relatively large bodies (e.g., ungulates, carnivores, and mesocarnivores) were substantially more common than those with small bodies (e.g., birds, reptiles, amphibians, and small mammals). These biases might reflect the high data requirements for many of the methods in movement ecology, meaning that only extensively monitored systems are regularly considered at the level of individual hosts. Alternatively, the taxonomic bias in hosts could be indicative of technological limitations that, until recently, prohibited the tracking of animals with smaller bodies with advanced instruments; alternatively, taxonomic bias patterns closely track phylogenetic hotspots of zoonotic and agriculturally-relevant pathogens.

**Figure 6:**
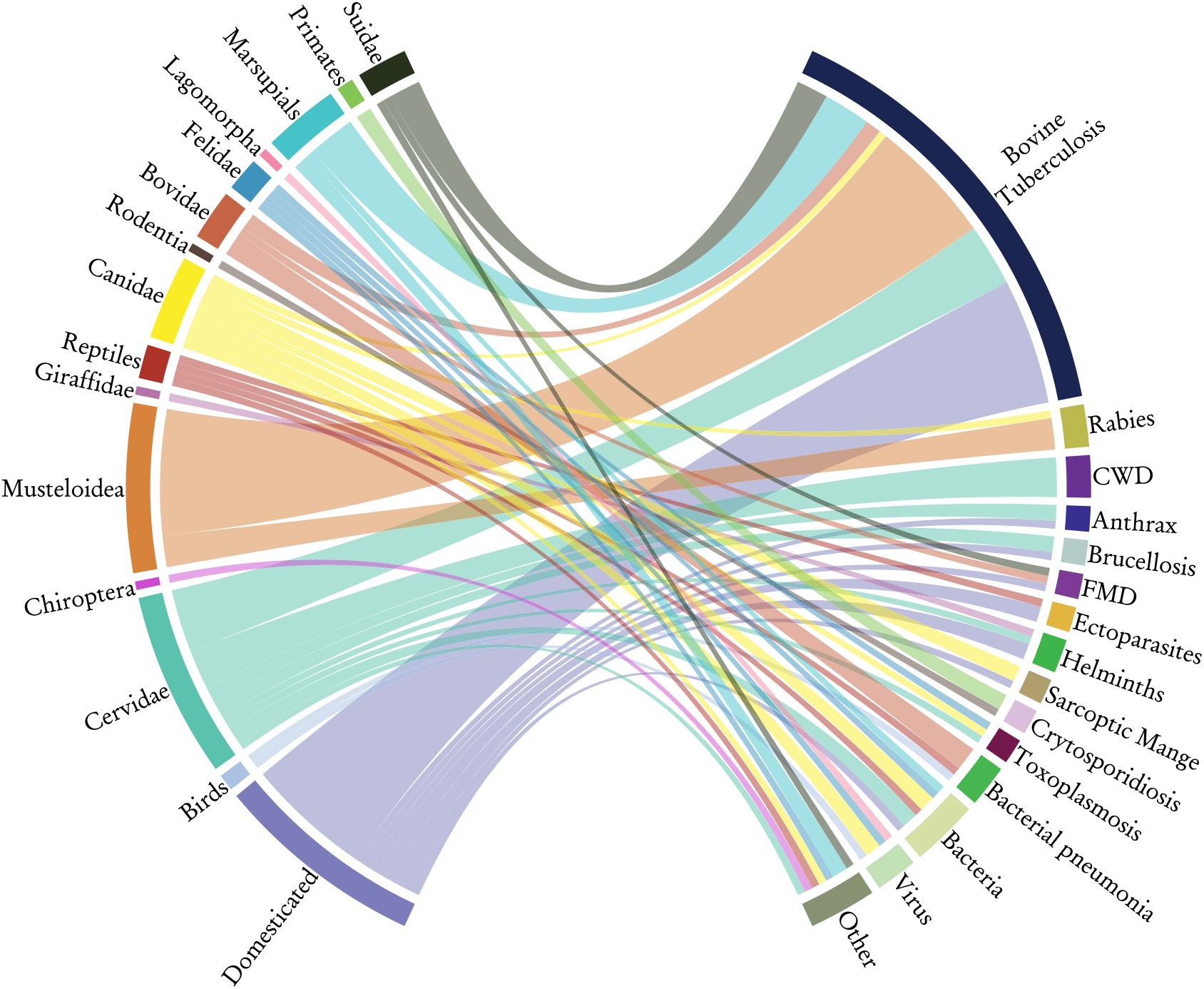
Study Bias in Movement & Disease Ecology Literature. We preformed a systematic review of scientific literature, identifying 70 studies currently using movement data and methods within disease research since 2000. In the above chord diagram, the host taxonomic order (left) is linked with the associated pathogen or parasite taxon (right), with the width of the bar indicating the proportion of studies investigating that particular pairing. Expectedly, pathogens with possible spillover threats to humans or livestock receive most of the attention. For example, studies of bovine tuberculosis (*Mycobacterium tuberculosis*) systems were particularly prevalent in the literature, likely because of the risk faced by cattle in proximity to possums, raccoons, badgers, and other mammals. Other well studied pairings included bighorn sheep with bacterial pneumonia (*Mycoplasma ovipneumoniae*); raccoons and canines with rabies; and deer with various livestock spillover diseases, such as anthrax (*Bacillus anthracis*), brucellosis (*Brucella abortus*), foot and mouth disease (FMD; *Aphthae epizooticae*), and chronic wasting disease (CWD).

For the 70 studies that met the criteria for inclusion, all methods of analyses used by the researchers were sorted into four broad groups: spatial overlap, habitat selection, network analyses, and behavioral analyses. In several cases, more than one of these methods were used in a single study, resulting in a total of 91 analyses. Spatial overlap was the most frequently used analysis, with 41 cases applying some form of overlap method. These ranged from examinations of home range dynamics (e.g., Yockney *et al.* 2013) to studies that attempted to measure the number of contacts between animals (e.g., Woodroffe & Donnelly 2011), often using proximity sensors to do so (e.g., Marsh *et al.* 2011). Habitat selection analyses were also quite common, with 24 cases using selection functions (e.g., Morris *et al.* 2016) or performing basic comparisons between habitat types (e.g., Parsons *et al.* 2014). Similarly, studies that drew upon the wide array of network analysis tools were fairly common, with 19 constructing some form of network, often with the use of proximity sensors (e.g., Hamede *et al.* 2009). The least common form of analyses encountered during the literature review were behavioral analyses, where researchers explicitly measured the probability of a particular behavior (e.g., dispersal; Caron *et al.* 2016) or compared individuals of two different behavioral classes (e.g., migratory vs. resident; Pruvot *et al.* 2016). Only 6 cases of behavioral analysis appeared in the resulting literature. Since the role of behavior in influencing disease dynamics is well established, this represents an underexplored avenue for investigation of disease systems.

There was a demonstrable correlation between the mode of transmission (Figure 6) exhibited by a pathogen and the methods ultimately selected to study it. Although some studies (13) did not identify a transmission mode, many emphasized that whether the pathogen studied had a direct (20) or indirect (11) transmission route. Many studies (26), mostly on bovine tuberculosis, mention that both transmission modes are possible, but researchers often selected their methods based on one or the other (4 of the 26 emphasize direct transmission, while 7 focus on indirect). Of those studies focused on the indirect mode of transmission, spatial overlap methods were used in approximately 56%, habitat selection in about 44%, network analyses in nearly 17%, and behavioral analyses in only 6%. By contrast, studies of direct transmission used network-based analyses (46%) and behavioral analyses (17%), but spatial overlap methods were nearly as common as in studies of indirect transmission (50%), and habitat selection methods were far less common (13%). These differences are to be expected: pathogens with particular transmission modes require the use of tools and methods relevant to the movement processes that underlie them.

## Discussion and Future Directions

Complex patterns in ecology frequently emerge from simple rules at fine scales. As we highlight, basic rules of animal behavior drive the complex interplay of animal movement and disease dynamics; the implications for wildlife and human health are major. Incorporating movement behavior into epidemiological models could improve predictions of disease dynamics, provided the additional level of complexity is handled correctly (Getz *et al.* 2017). While we have highlighted specific well-developed pairs of pathogen transmission mode and analysis methods (like networks and direct contact pathogens, or landscape models and vector-borne disease), we also note that many pathogens exploit several transmission strategies, and researchers will correspondingly need several methods in these cases. Developing protocols that include movement data in basic disease research, and vice versa, will be an important first step towards making these advances more feasible—and towards making broad advances in ecological theory, as some disease ecologists have begun to do with network methods (Sah *et al.* 2017a).

Movement tools will likely increase in value with ongoing improvements in biologging technologies (Kays *et al.* 2015). For example, advancements in radar and radio-frequency technologies allow tracking of a broader range of insect movements (Kissling *et al.* 2014), offering the potential to include these movements when considering vector-borne disease dynamics. Further, accelerometer-based data and very-high resolution GPS tracking (e.g., 1 Hz fix rates) will help researchers parse movement tracks at an even finer scale than current path segmentation methods allow (McClintock *et al.* 2017). In doing so, proximity-based social networks could be further informed with the behavioral states of individuals, potentially clarifying the epidemiological relevance of such points of contact (Spiegel *et al.* 2016; Sih *et al.* 2017; Spiegel *et al.* 2017b). The decreasing costs of these technologies could soon offer opportunities to monitor entire populations, thereby shifting researchers from extrapolating risk across a population to measuring contact rates directly. More comprehensive surveillance may also enable the development of models that more accurately infer dose exposure, based on duration of contact between animals and infected hosts or environmental reservoirs, vastly improving models of the heterogeneity in transmission efficiency.

Though the host-environment and host-pathogen interactions reflected in movement data can offer significant insight into disease dynamics, important processes might also occur at the pathogen-environment (or vector-environment) interface. In benthic marine systems, for example, suspension-feeders that filter large volumes of water while feeding can be particularly vulnerable to infection by microparasitic pathogens floating in the water (Lafferty 2017). This accumulation process has been modeled through the incorporation of particle diffusion (Bidegain *et al.* 2016), but the nature of these pathogens and their deposition makes the precise tracking of their movements in such dynamic environments very difficult. Thus, the validity of forecasts based on host movement alone is in question when pathogen-environment interactions (e.g., pathogen movement, rates of growth or decay, or the length of vector life history stages) occur at time scales comparable with the host-pathogen interactions themselves (e.g., lengths of latent and infectious periods). When response time scales are comparable, coupled host-pathogen-environment models are required. Though this has not been the emphasis of much of the recent work in movement ecology, the expansion of methods and technologies to accurately track minute particles through three-dimensional space is a frontier worthy of exploration. The resulting models could replace assumptions regarding the diffusion of such particles and further aid in our understanding of contact processes in highly dynamic environments.

Although we have focused on host populations, these tools also apply to multi-species transmission, such as in the spillover of wildlife diseases into livestock, or spillback of diseases from domesticated animals into wildlife (Barasona *et al.* 2014). Furthermore, these methods could just as easily be used to assess the risk of zoonotic spillover into human populations. Currently, ecological niche modeling is a popular proxy for zoonotic disease risk, but this only summarizes high-level landscape patterns (often treating host-pathogen systems as one coupled phenomenon); replacing these, or combining them, with movement models like RSFs can more accurately characterize average or seasonal patterns of host movement, and therefore risk to human health. In particular, in the case of pathogens that affect free-ranging and often migratory hosts such as bats (i.e., Ebola, Marburg, or Nipah viruses), overlap analyses could illuminate potential risk zones for future spillover events. With additional data collection using advanced monitoring devices, researchers can move beyond treating overlap (spatial or spatiotemporal depending on the pathogen or parasite in question) as a proxy for contact; in fact, we note the clear but unexplored potential for animal movement studies to act as part of a realtime early warning system for difficult-to-surveil zoonoses.

With a common language and mutual appreciation for their respective disciplines, disease ecologists and movement ecologists can collaborate to help solve pressing problems. Like Ebola or Nipah, most emerging diseases spill over from wildlife (Jones *et al.* 2008). Controlling such diseases is difficult, and interventions can be controversial (e.g. wildlife cullings), infeasible (e.g. mass wildlife or livestock vaccination), or ineffective; for example, culling badgers can spread bovine tuberculosis because badgers will move into treated areas (Woodroffe *et al.* 2006). Studying animal movement might help predict disease spread (and help explain why some interventions fail), and identify new interventions, such as wildlife relocations or vaccination. Furthermore, movement ecologists can benefit from considering how parasites alter animal movement, thereby accounting for otherwise unexplained variation in movement among individuals. Advances in disease diagnosis, combined with new technologies that and remotely monitor an animal’s physiology and motion make this an opportune time for studies to embrace both disease ecology and movement ecology.

## Acknowledgements

The authors would like to acknowledge the Getz and Brashares Labs for their extensive feedback. This research was supported by grants NIH GM083863 (WMG) and NIH GM117617 (PI Jason Blackburn; ED and DS supported under a UC Berkeley subcontract). We also appreciate the invaluable guidance offered by Sadie Ryan, Jason Blackburn, and Jane Flegal. We also thank Matthew Silk, Kevin Lafferty, and anonymous reviewers for their considerable and thoughtful feedback that greatly improved our revisions.

